# An oviposition pheromone, hexadecanoic acid, found on the eggs of *Phlebotomus papatasi* from Central Tunisia, attracts gravid females and stimulates oviposition

**DOI:** 10.1101/2025.05.01.651664

**Authors:** I. Chelbi, E. Zhioua, M. Shocket, J.G.C Hamilton

## Abstract

*Phlebotomus papatasi* is a vector of *Leishmania major*, the etiologic agent of zoonotic cutaneous leishmaniasis, a disfiguring and debilitating disease. In this study, we identified fatty acids found on the exterior of eggs laid by female *P. papatasi* that could be potential oviposition pheromones. We tested the effect of different treatments on 1) the number of eggs laid and 2) the spatial distribution of eggs laid. The treatments included three quantities of eggs (80, 160 and 320), hexane extracts of 160 eggs, 160 eggs after being washed with hexane to remove any pheromones, and three concentrations (1ng, 10ng and 100ng) of synthetic versions of three fatty acids that we identified as being present on egg exteriors.

The saturated fatty acids dodecanoic (C12) and tetradecanoic (C14) acid, identified by GC/MS analysis, were abundant in hexane extracts of both eggs and gravid females but were present in only trace amounts in males. Hexadecanoic and hexadecenoic (C16) acids were abundant on eggs, gravid females and males.

A negative binomial GLM found that significantly more eggs were oviposited by gravid females in response to 80 eggs (*P*=0.0255), 160 eggs (*P*<0.001), 320 eggs (*P*<0.001) and the hexane extract of 160 eggs (*P*<0.001). Eggs washed in hexane were not more attractive than a control (*P*=0.591). The number of eggs laid was increased by all three concentrations of hexadecanoic acid (*P*<0.001), 10ng and 100ng of tetradecanoic acid (*P*<0.001), and 1ng and 10ng of dodecanoic acid (*P*<0.001).

The spatial response of oviposition (the proportion of eggs laid on the test vs. control side of the oviposition pot) was weaker than the response of total eggs laid. A beta GLM found that gravid females laid a significantly higher proportion of eggs near 160 eggs (*P*=0.004) and significantly lower proportion of eggs near 100ng of dodecanoic acid (*P*=0.016). Bootstrapping and permutation tests also suggested significant attractive effects of 320 eggs, egg extract, and 1ng and 10 ng of hexadecenoic acid.

These results suggest that hexadecanoic acid is the oviposition pheromone, of *P. papatasi* from Tunisia because its presence increases both the number of eggs laid and attracts oviposition over the small spatial scales of the assay. Studies by others have shown that dodecanoic acid is the oviposition pheromone of *P. papatasi* from Turkey. In this study, dodecanoic acid increased the number of eggs laid but either did not change their spatial distribution or was repulsive at the highest concentration. The observed difference may be related to the different geographical origins of the sand flies used in this study.

**Author Summary:** Zoonotic cutaneous leishmaniasis, caused by the parasite *Leishmania major,* is a disfiguring and debilitating disease. It poses a significant public health problem worldwide, particularly in North Africa, the Middle East, and Eastern Europe. The parasite is transmitted from rodent reservoirs to human hosts through the blood-feeding activity of the sand fly vector, *Phlebotomus papatasi*. There is a lack of understanding of the basic elements of the ecology of this sand fly species, and in particular, the role of volatile chemicals in the identification and location of oviposition sites. In his study, we showed that hexadecanoic acid, a fatty acid present on the eggs of Tunisian *P. papatasi*, was attractive to other gravid (egg-laying) female sand flies and stimulated their egg laying. In addition, comparison of these results with previously published provide evidence for a hitherto unrecognised population sub-structuring in *P. papatasi* and a potential tool for control.

## Introduction

Female *Phlebotomus papatasi* Scopoli (Diptera: Psychodidae) sand flies are the main vector of *Leishmania major*, the etiologic agent of zoonotic cutaneous leishmaniasis (ZCL) in the Old World (Ben Ismail *et al*. 1987; Derbali *et al*. 2012). Cutaneous leishmaniasis, which affects about 1.5 million people per year, is considered by the WHO to be a neglected tropical disease (Alvar *et al*. 2012). The disease is widespread in the Old World from Morocco to Saudi Arabia to Afghanistan where it constitutes a serious public health problem (Alvar *et al*. 2012). It is often strongly correlated with poverty (Alvar *et al*. 2006) and although not fatal, the lesions produced may cause substantial disfigurement and severe distress with lifelong psychological and social consequences (Kassi *et al*. 2008).

ZCL affects thousands of people in the endemic areas of Central and Southern Tunisia, with an annual average incidence rate of 502.9 cases per 100,000 inhabitants (Bellali *et al*. 2015; Chelbi *et al*. 2007; Chelbi *et al*. 2009). As no available vaccine for ZCL exists, new vector control methods and strategies are urgently needed (Ben Hadj Ahmed *et al*. 2010).

In North Africa, the fat sand rat, *Psammomys obesus*, and Shaw’s jird, *Meriones shawi*, are the principal reservoir hosts of *Le. major* (Ben-Ismail *et al*. 1987; Fichet-Calvet *et al*. 2003; Ghawar *et al*. 2011; Rioux *et al*. 1992; Rioux *et al*. 1986). Both species inhabit complex burrow systems [El-Bana 2009] associated with their food sources; saltbush *Atriplex halimus* (Chenopodiaceae) and jujube tree *Ziziphus spp.* (Rhamnaceae) respectively (Derbali *et al*. 2013). The burrows have moderate, stable temperatures and elevated humidity which creates a suitable microclimate for the immature and adult stages of *P. papatasi* (Derbali *et al*. 2014). Adult female *P. papatasi* utilize the rodents as the source of their primary blood meal, and the rodent faeces and plant debris that accumulate in the burrows are the main food source for the sand fly larvae (Derbali *et al*. 2014). Rodent burrows are therefore considered to be the main breeding sites for *P. papatasi* (Derbali *et al*. 2014).

Developing vector control strategies based on the manipulation of chemically mediated behaviour remains an option for both New and Old-World sand flies (Hamilton 2022). For example, the application of a synthetic sex/aggregation pheromone in a lure-and-kill (attract-and-kill) approach for *Lutzomyia longipalpis*, the vector of visceral leishmaniasis (VL) in Central and South America, has been shown to reduce canine *Leishmania infantum* parasite load and infectiousness, and to reduce the numbers of sand flies significantly in the peri-domestic environment (Courtenay *et al*. 2019). If widely implemented, this control measure could significantly reduce human and canine VL cases (Retkute *et al*. 2021).

Gravid *Lu. longipalpis* are attracted to and stimulated to oviposit by dodecanoic acid, deposited on eggs during oviposition (Dougherty and Hamilton 1997; Elnaiem and Ward 1990; Elnaiem and Ward 1991; Elnaiem and Ward 1992) and kairomones from the oviposition substrate (Dougherty *et al*. 1993; Elnaiem and Ward 1992). It has been proposed that these chemical cues potentially improve progeny survival (Killick-Kendrick 1990). Similarly gravid *P. papatasi* have also been shown to be attracted to and to lay significantly more eggs by chemicals which are present on conspecific eggs (Kowacich *et al*. 2020; Srinivasan *et al*. 1995). Dodecanoic acid, found in both egg extracts and 1^st^ stage larvae of *P. papatasi* from Abkük, Turkey was identified as the oviposition pheromone (Kowacich *et al*. 2020). This pheromone and kairomones from the decaying oviposition substrate, bacteria, host animals and conspecific larvae are believed to guide gravid *P. papatasi* in choosing rodent burrows for oviposition (Faw *et al*. 2021; Kakumanu *et al*. 2021; Marayati *et al*. 2015).

*P. papatasi* has a very wide geographical distribution. It occurs from Morocco in the West across the Middle East to the Indian subcontinent and from southern Europe to north, central and eastern Africa in diverse ecological settings (Lewis 1982; Maroli *et al*. 2013). Although there have been suggestions that *P. papatasi* is a species complex based on genetic differences between the Tunisian and Turkish populations (Hamarsheh 2011; Hamarsheh *et al*. 2009), the weight of evidence is largely against that proposition (Depaquit *et al*. 2008; Ghosh *et al*. 1999).

Selective response to semiochemicals from food and other resources as well as the production and response to pheromones (e.g. sand fly sex/aggregation pheromones) can provide evidence of species isolation (Hamilton *et al*. 2005; Hickner *et al*. 2021; Labbé *et al*. 2023; Maingon *et al*. 2003). Although behavioural evidence suggests the presence of a sex pheromone in *P. papatasi,* no chemical or potential sources have been identified (Chelbi *et al*. 2012; Chelbi *et al*. 2011). Therefore, in this study we examined the presence and determined the structure of the oviposition pheromone of *P. papatasi* from Tunisia and considered its potential as a marker of population sub-structuring.

### Methods Sand flies

A colony of *P. papatasi e*stablished from adults originally collected in El Felta (34° 5’N, 9° 29’E), Sidi Bouzid, Tunisia in 2003 was maintained in the vector ecology laboratory at Institute Pasteur de Tunis (Chelbi and Zhioua 2007). Newly emerged *P. papatasi* were kept in Barraud cages (20 x 20 x 20cm) in an insectary (27°C; 80% rh; photoperiod 12:12 (L:D)). Three days after emergence, females were blood-fed on anaesthetized mice and then offered a 30% sucrose solution. Blood-fed females were removed from the feeding cage after 1h, placed with an equal number of males in a Barraud cage, and left for 4 days to allow complete oogenesis and defecation. After 5 days, gravid females were transferred into Nalgene plastic pots (BDH, Speke, Liverpool, UK), prepared by lining the base and walls with Plaster of Paris, to oviposit or were used in oviposition experiments.

### Chemical Analysis

Extracts of eggs, adult females and adult males were prepared for chemical analysis by placing batches of 1-day old eggs (n=1000), gravid females (n=15) or 3-4 day old males (n=30) in small clean screwcap glass reaction vials (2ml) and adding hexane (100µl). The vials were capped and left for 48h (-20°C) to allow complete extraction and to minimize loss of volatiles.

To convert fatty acids to their methyl esters for gas chromatography coupled mass spectrometry (GC/MS) analysis the extracts were transferred to cleaned glass screwcap reaction-vials (2ml) and reagent (100µl BF3-methanol) (Sigma-Aldrich) was added and heated (60°C) for 25 minutes. After cooling, hexane (1ml) was added and the solution washed two times with saturated NaCl (1ml). The remaining hexane layer was dried (anhydrous sodium sulphate) and transferred to a clean glass vial and the volume reduced under N2 to 100µl.

Identification of the fatty acid methyl esters was performed by gas chromatography coupled mass spectrometry (GC-MS) (Dougherty and Hamilton 1997). Briefly, analysis was carried out on a Hewlett Packard 5890II+ gas chromatograph with an HP-5MS column (Hewlett Packard UK; 30m x 0.25mm ID, 0.25µm film thickness) coupled to a Hewlett Packard 5972A mass spectrometer (EI, 70eV, 165°C). Samples were introduced via a split/splitless injector (splitless mode, sampling time 0.7min, 180°C). The HP- 5MS column temperature was initially held at 50°C (4min), then increased (5°C min^-1^) to 310°C (10min). The carrier gas was helium under constant pressure with carrier gas flow rate of 1ml min^-1^.

*Phlebotomus papatasi* egg (1µl, equivalent to 10 eggs), gravid female (1µl, equivalent to 0.15F) or male (1µl, equivalent to 0.3M) extracts were injected onto the HP-5MS column. The fatty acid methyl esters were identified by comparison with spectra held in the NIST spectral library (Dougherty and Hamilton 1997) and by retention time and mass spec comparison with standards prepared from the methyl ester of dodecanoic acid (C12) (DDA), tetradecanoic acid (C14) (TDA), hexadecanoic acid (C16) (HDA), 9- hexadecenoic acid, 7-hexadecenoic acid and 2-hexadecenoic acid (Sigma-Aldrich Ltd).

### Oviposition experiments general protocol

Round-tipped forceps were used to score the Plaster of Paris surface of oviposition pots to create twelve 1cm long, 3mm deep grooves arranged in two adjacent 2×2 grid patterns repeated on the opposite side of the pot (Figure 1). The grooves were created to induce thigmotrophic oviposition previously observed in other species of sand flies (Elnaiem and Ward 1990). One side of the oviposition pot was designated the test side and the other the control. Pots were moistened with distilled water and placed on tissue paper for half an hour to eliminate any excess water. Five-day-old gravid female *P. papatasi* (n=30) were then introduced into an oviposition pot which was then placed within a black bag placed inside an incubator (29-30°C, 95% rh) (Chelbi and Zhioua 2007). Oviposition was allowed to continue in complete darkness (which removed visual oviposition cues) for 4 full days. On the 5^th^ day, all eggs laid in the test and control sides of each pot were counted under a dissecting microscope.

**Figure 1.**
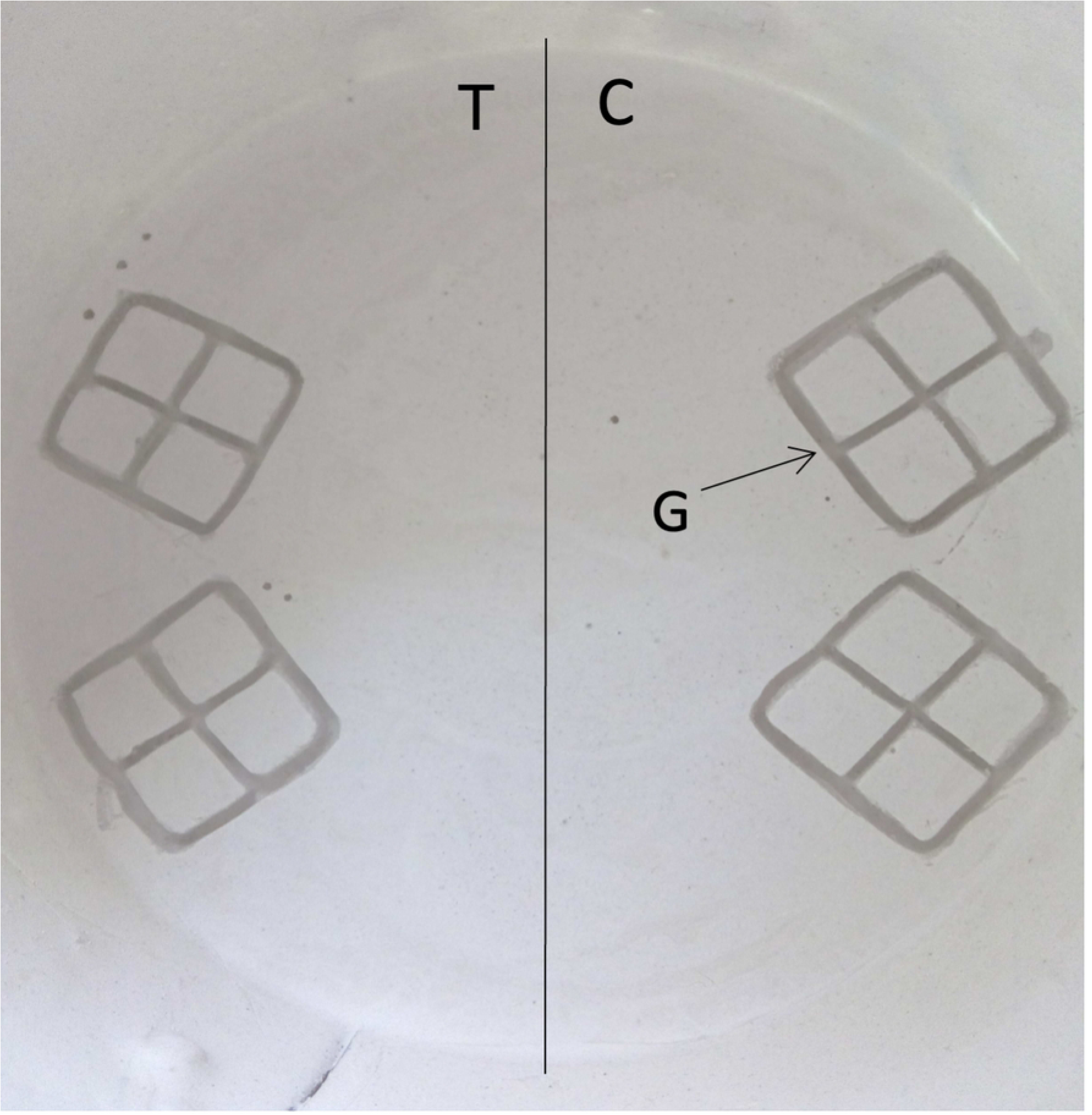
Plan view of a freshly prepared oviposition pot with grooves etched into the surface of the Plaster of Paris. One half of the oviposition pot containing two adjacent 2×2 grid patterns was designated the test side (T) and the other side the control (C). The grooves (G) were etched in the surface of the Plaster of Paris with round tipped forceps. Egg counting was done separately for the whole of the test side and the whole of the control side.

### Oviposition experimental treatments

We conducted a series of oviposition experiments with the following treatments. The number of replicates performed for each treatment varied from 6 to 10 for control trials, 9 to 12 for egg trials, and 14 to 16 for synthetic fatty acids trials (Table 2).

### Two-blank control

To measure the baseline number of eggs and the variance in the spatial distribution of eggs in the absence eggs, extract of eggs, or synthetic fatty acids, both sides of the oviposition pot were left untreated (blank vs blank).

### Hexane

To determine if the hexane solvent stimulated or directed oviposition, we placed hexane (10µl) on a piece of filter paper (1cm diam.) on the test side of the oviposition pot between the two adjacent 2×2 grid patterns (Fig. 1). On the control side filter paper with nothing added (hexane vs blank).

### Washed eggs

To determine if the physical presence of eggs alone (with the surface chemicals removed) stimulated or directed oviposition, we placed 160 eggs that had been washed in hexane for 48h in the test side of an oviposition pot and nothing in the control side (washed eggs vs blank).

### Unmodified eggs

To determine whether previously laid eggs stimulated or directed oviposition, freshly laid (1-day-old) *P. papatasi* eggs (either 80, 160, or 320) were transferred with a dissecting needle into the grooves on the test side of the oviposition pot. The control side was left untreated (eggs vs blank).

### Egg extracts

To determine if hexane extracts of freshly laid eggs stimulated or directed oviposition, extracts were prepared by placing 160 eggs in 10µl of hexane for 48h (-20°C). Afterwards, the extract was removed and placed on a piece of filter paper (2cm diam) placed in the ‘test’ side of the oviposition pot. The same volume of hexane only (10µl) was also placed on a piece of filter paper (2cm diam) in the control side of the pot, directly opposite the test side (egg extract vs hexane).

### Synthetic fatty acids

To determine if synthetic copies of the fatty acids (FAs) identified by GC/MS in eggs and gravid females stimulated or directed oviposition, three quantities (1ng, 10ng or 100ng in 10ul hexane) of synthetic FAs (dodecanoic acid [DDA], tetradecanoic acid [TDA] and hexadecanoic acid [HDA]) (Sigma Aldrich UK) were placed (individually) on filter paper in the test side of the oviposition pots. The same volume of hexane only (10µl) was placed on a piece of filter paper in the control side of the pot directly opposite the test side (synthetic fatty acids vs hexane). We also tested the oviposition response to two fatty acids structurally similar to 9-hexadecenoic acid: 7- hexadecanoic acid and 2-hexadecenoic acid (9 times each).

### Statistical Analysis

We analysed the data with generalised linear models (GLMs) fit in R (R Core Team 2023) . The count data for total eggs laid were over-dispersed for a Poisson GLM, so we modelled them with a negative binomial GLM using the “MASS” package (Venables and Ripley 2002). We modelled the spatial preference for egg laying as the expected proportion of eggs laid on the test side versus the control side of the oviposition pot. We fitted a beta GLM using the “betareg” package (Cribari-Neto and Zeileis 2010). For both GLMs, we fitted one model with data from all the oviposition trials and used the two- blank control treatment as the baseline (intercept for contrasting with the other treatments).

We conducted two additional analyses on the proportion of eggs laid on the test side of the oviposition pot. First, we bootstrapped 95% confidence intervals for the mean proportion of eggs laid on the test side for each treatment. Specifically, we sampled with replacement from the dataset, where the number of samples matched the original number of replicates for a given treatment, with 1000 bootstrapped iterations. Second, we performed permutation tests, where the “test side” and “control side” labels were assigned to trials in every possible permutation to generate a probability distribution for each treatment. We then calculated the cumulative probability density for the observed mean proportion of eggs on the test side for each treatment. The bootstrapped CIs and permutation tests allowed us to test the null assumption that there is no spatial preference and the mean proportion of eggs on the test side = 0.5.

## Results

### Chemical Analysis

Chemical analysis of the derivatised *P. papatasi* egg extract revealed the presence of fatty acid methyl esters in the hexane extracts of 1000 eggs, 15 gravid female and 30 male *P. papatasi*. The main FA components present were; HDA, TDA and DDA. DDA and TDA were significant components of the eggs and gravid female extracts but were present in trace amounts (<1%) in males HDA was present in extracts of all 3 sources in similar proportions Table 1. The unsaturated FA, 9-hexadecenoic acid, was tentatively identified in extracts from all 3 sources in similar proportions. Octadecanoic acid (ODA) was present in trace amounts in males and absent from gravid females and eggs. The tentatively identified unsaturated FA, 9,12-octadecadienoic acid, was absent from both eggs and gravid females but was present in males. In addition, 9-octadecenoic acid was tentatively identified as present in small amounts in eggs and gravid females, but it was abundant in males.

**Table 1.** Abundance of main fatty acids present in extracts of *P. papatasi* eggs, gravid females and males.

| fatty acid | molecular weight | abundance (%) |  |  |
| --- | --- | --- | --- | --- |
|  |  | eggs | gravid females | males |
| dodecanoic | $C_{12}H_{24}O_2$ | 37.4 | 43.1 | 0.3 |
| DDA | 200.3 g mol<sup>-1</sup> |  |  |  |
| tetradecanoic | $C_{14}H_{28}O_2$ | 19.0 | 19.3 | 0.8 |
| TDA | 228.4 g mol<sup>-1</sup> |  |  |  |
| hexadecanoic | $C_{16}H_{32}O_2$ | 25.1 | 16.7 | 27.7 |
| HDA | 256.4 g mol<sup>-1</sup> |  |  |  |

Extracts were prepared from batches of either freshly laid eggs (n=1000), gravid females (n=15) or 3-4 day old males (n=30). % abundance is of FAMES (amounts of contaminants and other minor/trace compounds are not included).

**Table 2:** Total eggs oviposited by treatment: number of replicates, mean, median, standard deviation, and *P*-values from the negative binomial GLM.

| *Treatment* | *Rep* | *Mean eggs* | *Median eggs* | *Std. dev* | *GLM P-value* |
| --- | --- | --- | --- | --- | --- |
| Two-blank Control | 6 | 155.7 | 138 | 48.0 | NA |
| Hexane Control | 10 | 149.3 | 138.5 | 113.3 | 0.867 |
| Washed eggs (160) | 10 | 178.0 | 168.5 | 71.5 | 0.591 |
| 80 eggs | 9 | 274.7 | 179 | 282.4 | 0.0255 |
| **160 eggs** | **12** | **489.7** | **475** | **156.1** | **<0.001** |
| **320 eggs** | **10** | **476.8** | **463** | **169.1** | **<0.001** |
| **Hexane extract of eggs (160)** | **12** | **479.2** | **452** | **141.4** | **<0.001** |
| **DDA – 1ng** | **15** | **567.5** | **549** | **222.7** | **<0.001** |
| **DDA – 10ng** | **15** | **527.0** | **608** | **231.3** | **<0.001** |
| DDA – 100ng | 15 | 214.5 | 195 | 112.6 | 0.170 |
| TDA – 1ng | 16 | 212.9 | 224.5 | 90.5 | 0.175 |
| **TDA – 10ng** | **14** | **593.5** | **708.5** | **323.5** | **<0.001** |
| **TDA – 100ng** | **15** | **477.6** | **452** | **1444.1** | **<0.001** |
| **HDA – 1ng** | **14** | **457.4** | **452** | **104.7** | **<0.001** |
| **HDA – 10ng** | **15** | **389.2** | **405** | **171.2** | **<0.001** |
| **HDA – 100ng** | **15** | **522.0** | **543** | **175.8** | **<0.001** |

DDA is Dodecanoic acid (C12), TDA is tetradecanoic acid (C14), HAD is hexadecanoic acid (C16). Rep is the number of replicates of each oviposition experiment. Mean eggs is the mean number of eggs laid in the oviposition pot. Std dev is calculated from all replicates. $P$ values $< 0.05$ are significant. Treatments that gave a significant result are in bold.

### Oviposition Experiments: total eggs

The total number of eggs laid increased significantly in response to the presence of untreated eggs, the hexane extract of eggs, and most concentrations of most of the synthetic fatty acids (Table 2, Figure 2). The two-blank control treatment established the baseline number of eggs oviposited (mean = 155.6 eggs, median = 138 eggs). The negative binomial GLM found no significant impact of hexane alone, 100ng of DDA, 1ng of TDA, or eggs washed with hexane (*P* > 0.1). All other treatments significantly stimulated oviposition, including all three quantities of unmodified eggs (80, 160, 320), hexane extract of 160 eggs, two quantities of DDA (1ng and 10ng), two quantities of TDA (10ng and 100 ng), and all three quantities of HDA (1ng, 10ng, and 100ng) (*P* < 0.05). On average, the significant treatments increased the mean number of eggs laid by 207% (mean of means = 477.7 eggs) and the median number of eggs laid by 248% (mean of medians = 480.6 eggs). The number of replicates, means, medians, standard deviations, and P-values from the GLM for specific treatments are provided in Table 2.

**Figure 2.**
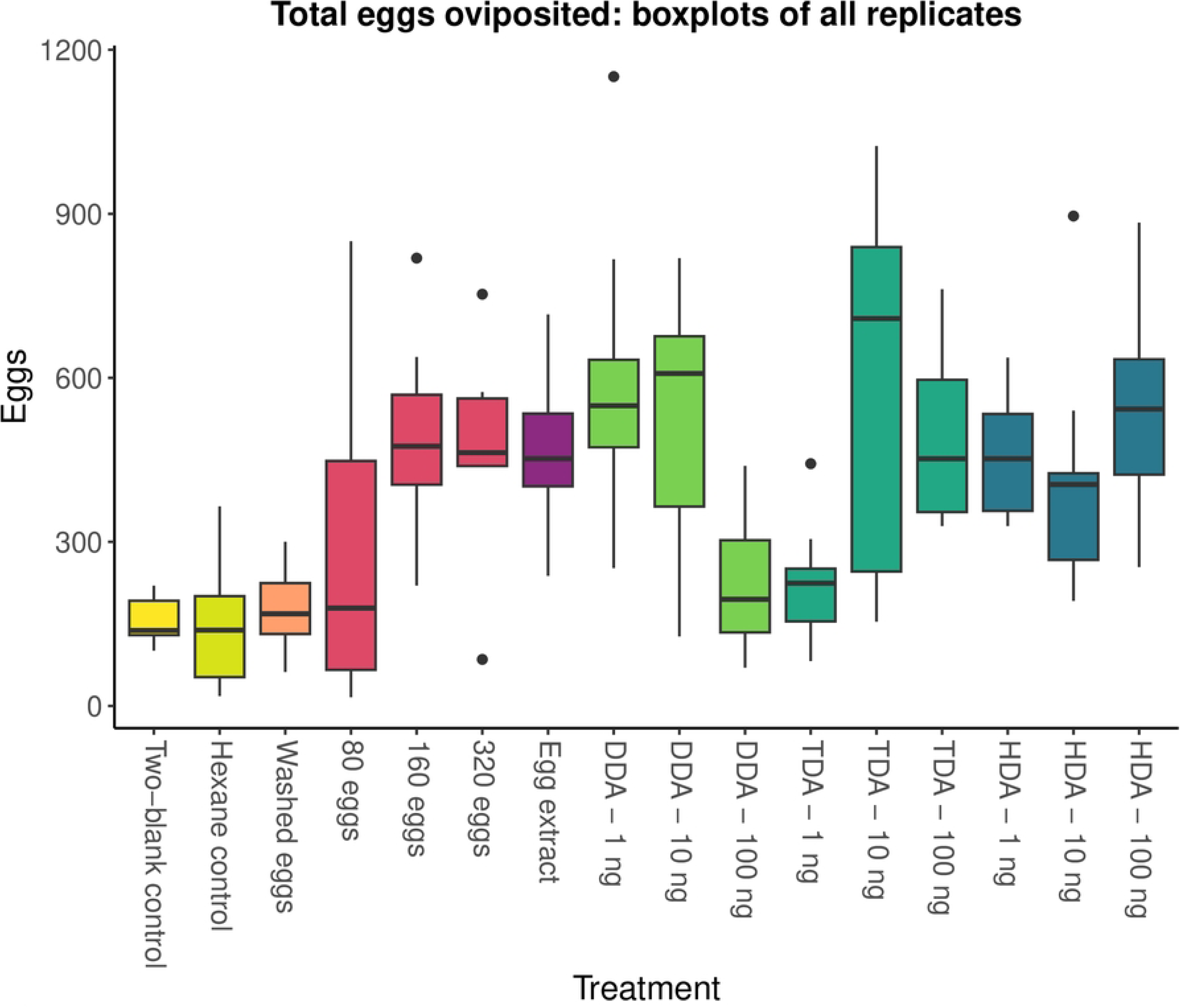
Boxplots of total eggs oviposited by gravid female *P. papatasi* in response to treatments of eggs and synthetic fatty acids. A negative binomial GLM found that all treatments are significantly different from the two-blank control (*P* < 0.05), except the hexane control, washed eggs, 100ng of dodecanoic acid (DDA), and 1ng of tetradecanoic acid (TDA).

### Oviposition Experiments: spatial preference and attraction

The effect of test material (eggs, extract and synthetic chemicals) on where eggs were oviposited (spatial preference) was weaker than the effect on the total number of eggs laid (Tables 3 and 4, Figure 3). The two-blank control treatment established the expected variance in oviposition location due to random processes. These data were highly variable: while half of the trials yielded results with eggs split relatively evenly between the two sides (52%, 53%, and 55% on the side with more eggs), the other half were extremely weighted to one side (67%, 70%, and 82% of the eggs on one side) and may be reflective of the oviposition process, i.e. once some eggs were laid then that side of the oviposition pot was favoured for further oviposition. This high level of variation limited our ability to detect significant differences in spatial preference in our experimental treatments with the beta GLM. Based on the beta GLM, two treatments significantly impacted the spatial preference for where eggs were oviposited: 160 eggs had an attractive effect (*P* = 0.004) while 100ng of DDA had a repulsive effect (*P* = 0.016). The numbers of replicates, means, medians, standard deviations, and *P*-values from the GLM for specific treatments are provided in Table 3.

**Figure 3.**
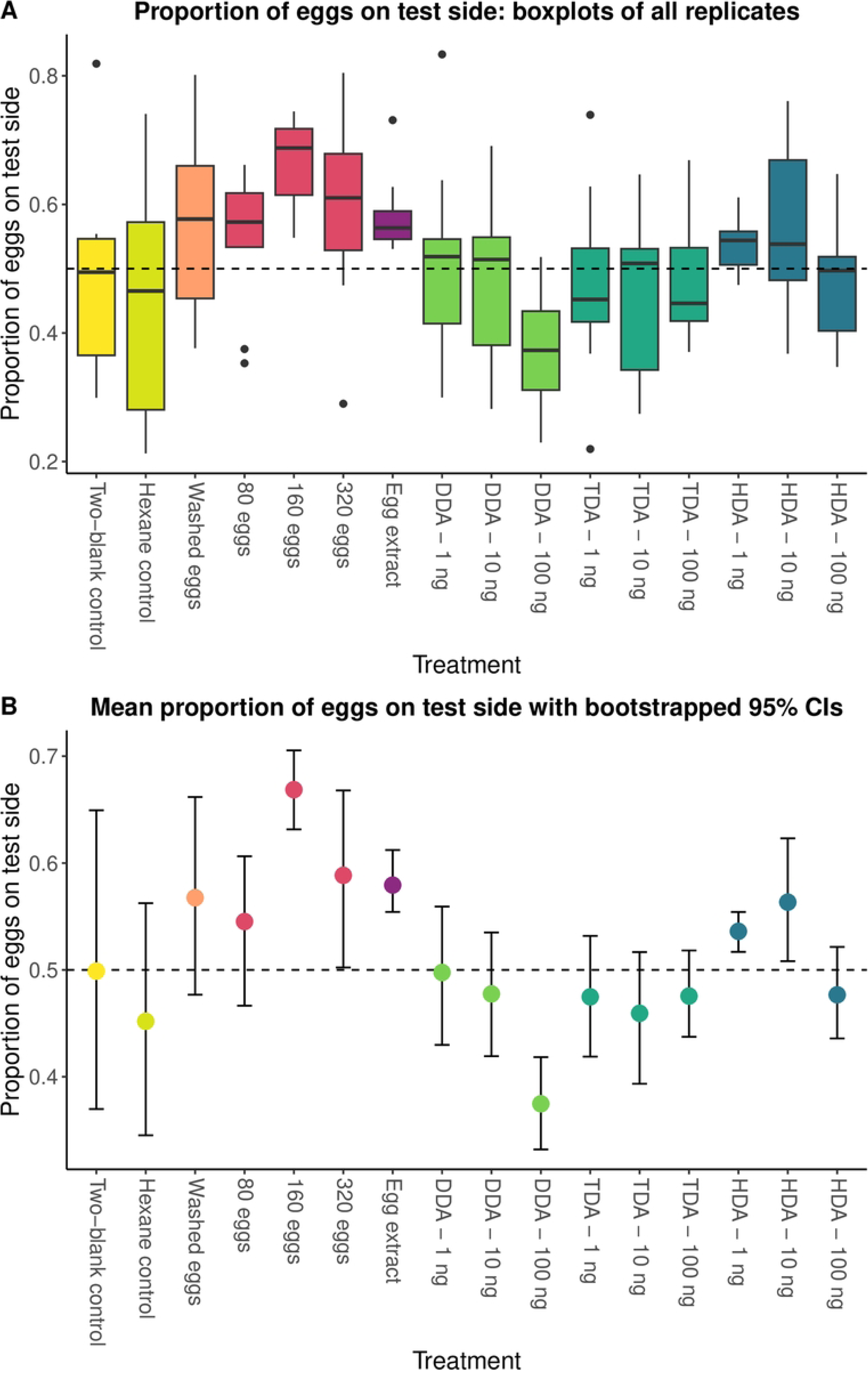
The spatial preference for oviposition by gravid female *P. papatasi* in response to treatments of eggs and synthetic fatty acids, measured as the proportion eggs laid on the test side of oviposition pots. Boxplots of the beta GLM (A), 160 eggs were attractive (*P* = 0.004), while 100ng of DDA was repulsive (*P* = 0.016) and boxplots of the mean proportion with bootstrapped 95% CIs and permutation tests (B) which were more sensitive. Eggs (n=320), egg extract, HDA (1ng and 10ng) were also attractive and significantly deviated from the null assumption that the mean proportion of eggs on the test side was = 0.5. Dashed lines in A and B mark where the proportion of eggs in either side = 0.5.

**Table 3:** Proportion of eggs laid on the test compared to control side of the oviposition pot by treatment: number of replicates, mean, median, standard deviation, and *P*-values from the beta GLM.

| *Treatment* | *Rep.* | *Mean prop.* | *Median prop.* | *Std. dev.* | *GLM P-value* |
| --- | --- | --- | --- | --- | --- |
| Two-blank Control | 6 | 0.500 | 0.495 | 0.187 | NA |
| Hexane Control | 10 | 0.452 | 0.465 | 0.181 | 0.322 |
| Washed eggs (160) | 10 | 0.567 | 0.577 | 0.153 | 0.243 |
| 80 eggs | 9 | 0.545 | 0.572 | 0.110 | 0.512 |
| **160 eggs** | **12** | **0.669** | **0.688** | **0.068** | **0.004** |
| 320 eggs | 10 | 0.588 | 0.610 | 0.143 | 0.141 |
| Hexane extract of eggs (160) | 12 | 0.580 | 0.563 | 0.055 | 0.196 |
| DDA – 1ng | 15 | 0.498 | 0.519 | 0.138 | 0.932 |
| DDA – 10ng | 15 | 0.478 | 0.514 | 0.116 | 0.607 |
| **DDA – 100ng** | **15** | **0.375** | **0.373** | **0.087** | **0.016** |
| TDA – 1ng | 16 | 0.475 | 0.452 | 0.119 | 0.571 |
| TDA – 10ng | 14 | 0.459 | 0.508 | 0.121 | 0.394 |
| TDA – 100ng | 15 | 0.476 | 0.446 | 0.087 | 0.597 |
| HDA – 1ng | 14 | 0.536 | 0.544 | 0.037 | 0.590 |
| HDA – 10ng | 15 | 0.563 | 0.538 | 0.121 | 0.265 |
| HDA – 100ng | 15 | 0.477 | 0.497 | 0.086 | 0.602 |

DDA is Dodecanoic acid (C12), TDA is tetradecanoic acid (C14), HDA is hexadecanoic acid (C16). Rep is the number of replicates of each oviposition experiment. Std dev is calculated from all replicates. GLM *P* values < 0.05 are significant. Treatments that gave a significant result are in bold.

**Table 4:** Results from bootstrapping and one-tailed permutation tests on the mean proportion of eggs laid on the test side of the oviposition pot: number of replicates, bootstrapped 95% CIs for the mean, and the cumulative density from the probability distribution of the permuted means that is greater than the observed mean proportion of eggs. Bolded treatments have 95% CIs that do not overlap 0.5 and probability density > 0.95 or < 0.05.

| *Treatment* | *Rep.* | *Bootstrapped*<br>*95% CI for mean*<br>*prop.* | *Permutation test:*<br>*probability density <*<br>*obs. mean prop.* |
| --- | --- | --- | --- |
| Two-blank Control | 6 | 0.375 – 0.644 | 0.500 |
| Hexane Control | 10 | 0.451 – 0.559 | 0.204 |
| Washed eggs (160) | 10 | 0.478 – 0.658 | 0.900 |
| 80 eggs | 9 | 0.469 – 0.606 | 0.883 |
| **160 eggs** | **12** | **0.633 – 0.702** | **1** |
| **320 eggs** | **10** | **0.503 – 0.665** | **0.959** |
| **Hexane extract of**<br>**eggs (160)** | **12** | **0.555 – 0.612** | **1** |
| DDA – 1ng | 15 | 0.433 – 0.561 | 0.474 |
| DDA – 10ng | 15 | 0.424 – 0.534 | 0.232 |
| **DDA – 100ng** | **15** | **0.331 – 0.419** | **9x10<sup>-5</sup>** |
| TDA – 1ng | 16 | 0.418 – 0.535 | 0.207 |
| TDA – 10ng | 14 | 0.400 – 0.518 | 0.113 |
| TDA – 100ng | 15 | 0.436 – 0.522 | 0.144 |
| **HDA – 1ng** | **14** | **0.518 – 0.554** | **0.998** |
| **HDA – 10ng** | **15** | **0.508 – 0.627** | **0.969** |
| HDA – 100ng | 15 | 0.437 – 0.517 | 0.156 |

DDA Dodecanoic acid (C12), TDA tetradecanoic acid (C14), HDA hexadecanoic acid (C16). Rep is the number of replicates of each oviposition experiment

Bootstrapped 95% CIs and permutation tests were more sensitive than the beta GLM, as they did not rely on comparison with the highly variable two-blank control and instead compared the observed proportion of eggs to the null expectation that the mean proportion of eggs on the test side would be 0.5. The bootstrapped 95% CIs and permutation tests found that 320 eggs, egg extract, 1ng of HDA, and 10ng of HDA also attracted oviposition and significantly deviated from the null expectation (Table 4).

## Discussion

Overall, these results established that oviposition response; attraction and stimulation, were to hexadecanoic acid, which could be isolated from the eggs of *P. papatasi*, and not the other compounds which were also present on the eggs.

Female *P. papatasi* were attracted to oviposit near freshly laid eggs. They were also stimulated to oviposit in the vicinity of freshly laid eggs. The response of gravid females to freshly laid eggs could be removed by washing the eggs in hexane. Eggs that had been washed with solvent failed to attract gravid females or to stimulate oviposition, indicating that the oviposition pheromone was present in or on the surface of freshly laid eggs. When the hexane extract of freshly laid eggs was placed on filter paper, the previously seen oviposition activity was observed, indicating that the pheromone could be transferred from the eggs to filter paper. Rescue experiments, to determine if the presence of cleaned eggs (in the presence of either egg extract or synthetic FAs on filter paper) provided tactile or visual stimulation, were not done. As the oviposition experiments were carried out in complete darkness visual cues are unlikely to play a significant role in oviposition however it remains possible that the eggs themselves provide tactile stimulation for oviposition. *Lutzomyia longipalpis* are known to be able to detect the physical properties of the oviposition substrate which is an important oviposition stimulus (Dougherty *et al*. 1993) and gravid females prefer to oviposit in vertical crevices rather than open, flat surfaces (Elnaiem and Ward 1992).

Fatty acids were identified in the egg extracts by coupled GC/MS analysis. The negative dose-dependent oviposition behaviour, previously observed in response to freshly laid eggs, was also seen with hexadecanoic acid (C16H32O2). HDA was significantly attractive at both 1ng and 10ng but was not attractive at 100ng, establishing the role of HDA as the key compound in directing oviposition. The other predominant (tetradecanoic and dodecanoic acid) and tentatively identified minor fatty acids (9-hexadecenoic acid, 7-hexadecenoic acid and 2-hexadecenoic) were not attractive and dodecanoic acid was repellent at the highest concentration (100ng) tested.

We did not test the response of gravid females to combinations of the fatty acids, however, the number of eggs laid in response to the egg extract was not substantially greater than the number laid in response to hexadecanoic acid.

For this study, we defined oviposition “stimulation” to mean the ability of eggs, egg extract or synthetic FAs to increase the number of eggs oviposited generally in the control side of the oviposition pot. This might also be described as “activation” as the females lay more eggs compared to the blank *vs* blank control. Eggs that were laid in the test side of the pot were considered to have been laid because of attraction. However, because of the proximity of the test and control sides as well as the potential role of freshly oviposited eggs to enhance the effect of the synthetic chemicals, extracts or experimentally placed eggs over the 4 days of the oviposition experiments the distinction between attraction and stimulation/activation is not clear cut.

Nevertheless, the observed changes in the number of eggs oviposited allow us to gain insight into the biological activity of the eggs, egg extracts and synthetic chemicals, on the numbers of eggs oviposited. Oviposition stimulation/activation occurred with all concentrations of HDA but only occurred when 100 or 10 ng of TDA and 10 or 1 ng of DDA were present.

Stimulation/activation did not occur with 1 ng of TDA or 100 ng of DDA. Further experiments with shorter assays (24 hrs) might reveal greater differences in the effect of the test material by reducing their potential spill- over effect. It is also worth considering that future experiments to further investigate the potential of HDA and other FAs to direct oviposition should more clearly separate potential attraction and stimulation/activation behaviour. This could be done by using olfactometer-based bioassays as well as field-based experiments.

We did not examine the response of gravid females to octadecanoic acid as both this chemical and the tentatively identified di-unsaturated octadecadienoic fatty acid were absent from both egg and gravid female extracts. Similarly, we did not examine the response of gravid females to the tentatively identified 9-octadecenoic acid which was abundant in males but present in small amounts in eggs and gravid females.

In this study, the number of eggs was important in determining the oviposition response, which was only significant when the number of eggs present was greater than 80. Previously, it was shown that *P. papatasi* from India laid significantly more eggs when 100 and 200 conspecific eggs were already present, but not in response to batches of 10, 20 or 40 eggs (Srinivasan *et al*. 1995). This phenomenon was also observed in a member of the New World sand fly species complex *Lu. longipalpis* (*s.l.*) where the presence of conspecific eggs (n≥80) resulted in increased oviposition (Elnaiem and Ward 1990). Hexane extracts of 1000 eggs significantly attracted gravid female *Lu. renei* in an olfactometer experiment over 30cm, whereas the extract of 100 or 200 eggs did not (Alves *et al*. 2003).

A previous study reported that the age of the eggs did not affect the number of eggs laid (Srinivasan *et al*. 1995), and in this study, we used 1-day-old eggs. However, it would be interesting to determine if the age of the flies was important in regulating their attraction to and stimulation by the pheromone.

Dodecanoic acid (C12) identified in the egg extracts of *P. papatasi* was shown to be attractive and stimulate egg laying in gravid females (Kowacich *et al*. 2020) and to be attractive to gravid *Lu. renei* (Alves *et al*. 2003). It has also been identified as the oviposition pheromone of *Lu. longipalpis*, the New World vector of *Le. infantum* (Dougherty and Hamilton 1997). Dodecanoic acid, the only saturated long-chain fatty acid present in the egg extracts, was found along with isovaleric acid (3-methylbutanoic acid). Hexadecanoic and tetradecanoic acid, found in this study were not reported in the egg extracts of *P. papatasi* examined previously (Kowacich *et al*. 2020).

The differences in chemical composition and biological activity seen between this study and that of Kowacich et al (2020) are substantial. Although the methodologies used were similar to each other they still had significant differences e.g. in our methodology the egg extracts were prepared directly from freshly laid eggs collected from the oviposition pot and placed in an extraction vial to which hexane solvent was added. In the work described by Kowacich et al, 2020 the eggs were rinsed off the plaster of Paris oviposition substrate with distilled water along with any dead flies, sieved to remove any debris and then left to dry on filter paper for 30 minutes. The eggs were then left for 12 h at 27 °C to reconstitute compounds that may have been removed during the egg collection before analysis. Depending on the quantity of water used and duration of immersion this might have affected FA content of extracts. In a further substantial difference, we did not find isovaleric acid (C5H10O2) in the egg extracts, whereas in the studies carried out by Kowacich et al 2020 it was the second most abundant compound.

It should be noted that the *P. papatasi* used in the Kowacich *et al*. (2020) study originated from Abkük, Turkey whereas the *P. papatasi* used in this study originated in Sidi Bouzid, Southern Tunisia. *Phlebotomus papatasi* has a very wide geographical distribution and occurs from Morocco in the West across the Middle East to the Indian subcontinent in the East and from southern European countries to north, central and eastern Africa in a diverse range of ecological settings (Lewis 1982; Maroli *et al*. 2013). Although there is little evidence to suggest that *P. papatasi* is a species complex (Depaquit *et al*. 2008; Ghosh *et al*. 1999), the subpopulation of *P. papatasi* in North Africa has been shown to be different from the Turkish subpopulation (Hamarsheh 2011; Hamarsheh *et al*. 2009). In addition, the Tunisian strain of *P. papatasi* is autogenous and the Turkish strain is anautogenous (Chelbi and Zhioua 2007) and other significant morphological and anatomical differences between *P. papatasi* populations in India have been noted (Srinivasan and Jambulingam 2012). It is possible therefore that the difference between our results and those presented by Kowacich et al. (2020) may be related to differences in aspects of the chemical ecology of *P. papatasi* in different parts of its range and are potential evidence of population sub structuring.

In *Lu. longipalpis* the oviposition pheromone, dodecanoic acid, is present in the accessory glands (Dougherty and Hamilton 1997; Dougherty *et al*. 1994; Dougherty *et al*. 1992). Female *Lu. longipalpis* obtain hexadecanoic acid from their blood meal and for 3 days convert it to dodecanoic acid (77%) and tetradecanoic acid (7%). In this study the rate of conversion of hexadecanoic acid to dodecanoic acid was much slower in *P. papatasi* with dodecanoic acid comprising ca. 43% of the FA content 5 days post blood feeding with dodecanoic acid accounting for ca. 43% of total FA abundance.

Several studies have shown that rabbit faeces, rabbit food, colony frass, and larval rearing medium all contain compounds that attract and stimulate oviposition in gravid *P. papatasi* (Schlein *et al*. 1990; Wasserberg and Rowton 2011), *Lu. longipalpis* (Dougherty *et al*. 1993; Elnaiem and Ward 1992; Peterkova-Koci *et al*. 2012; Wasserberg and Rowton 2011) and *Phlebotomus argentipes* (Kumar *et al*. 2013). In the future it will be important to determine the interaction between the oviposition pheromone and oviposition kairomones and the ability of gravid females to select oviposition sites that would be beneficial for their offspring.

In the future it will also be important to determine if hexadecanoic acid alone or in combination with the kairomones is attractive over distance and simulates oviposition in the field. Long chain fatty acids have in general low volatility and hexadecanoic acid by itself is not likely to be attractive over distance. The oviposition pot assays that we used in this study did not establish long range attractiveness. It will be important to establish which if any of the compounds present in the eggs are attractive over distance.

## Ethics Statement

Sand fly blood feeding at Keele University for colony maintenance was performed according to the guidelines and regulations of the Animals in Science Regulation Unit (ASRU) and in accordance with the terms of a regulated licence (PPL 40/3279) in compliance with the UK Home Office, Animals (Scientific Procedures) Act (ASPA) regulations. All procedures involving animals were reviewed and approved by the Animal Welfare and Ethical Review Board (AWERB) at Keele University.

Sand fly blood feeding at the Institut Pasteur de Tunis was performed under the Assurance of the US Office of Laboratory Animal Welfare [Assurance approval F-16-00170 (A5743-01)]. The Institut Pasteur de Tunis complies with the European Directive for the Protection of Vertebrate Animals used for experimental and other scientific purposes (2010/63/EU).

## Acknowledgements

We are grateful to Dr. Pam Taylor (Keele University) and Saifeddine Cherni (Institut Pasteur de Tunis) for assistance with the sand fly colonies. We are also extremely grateful to Professors Gideon Wasserberg and Coby Schal (North Carolina University Greensboro) who graciously commented in-depth, on an earlier version of the manuscript.

